# Distinct thresholds condition sense PTGS initiation and amplification

**DOI:** 10.64898/2026.05.05.722629

**Authors:** Martin Lacroix, Nicolas Butel, Andana Barrios, Agnès Yu, Nathalie Bouteiller, Ivan Le Masson, Hervé Vaucheret

## Abstract

It has long been known that *Pro35S-*driven sense transgenes have a high propensity to undergo post-transcriptional gene silencing (S-PTGS). However, what exactly conditions S-PTGS initiation and amplification to make it systemic remains unknown. Through genetic screens, we show that antagonistic chromatin-related mutations enhancing and reducing transgene expression, result in enhanced and reduced S-PTGS amplification capacities, respectively, without affecting the initiation rate. Analysis of a large set of independent transgenic plants confirm a direct relationship between transgene expression and its capacity to amplify S-PTGS. Combining an inducible or a tissue-specifically expressed *GUS* transgene with a *Pro35S:GUS* transgene locus prone to amplify S-PTGS but unable to spontaneously initiate it induces systemic S-PTGS, indicating that transient and/or local passing of a discrete threshold is sufficient to initiate S-PTGS. Together, these results call for the existence of distinct thresholds related to transiently produced aberrant RNA and permanently produced target mRNA levels, which condition S-PTGS initiation and amplification, respectively. We show that this model also applies to endogenous genes for which RNA Quality Control (RQC) acts as a first layer of protection against S-PTGS, and DCL2’s obscuration by DCL4 as a second layer, allowing RQC to dysfunction locally without translating into the drama of systemic S-PTGS.

## Introduction

Post-transcriptional gene silencing (PTGS) is an RNA-based mechanism acting primarily as a defense mechanism against invasive sequences such as viruses and transgenes, allowing plants to recover from virus infection under certain circumstances (Wingard 1928; Baulcombe 2004). The pathway is initiated by double-stranded RNAs (dsRNAs), which are processed by the DICER-like proteins DCL4 and DCL2 into short dsRNA duplexes of 21- and 22-nt, respectively, which are referred to as short interfering RNAs (siRNAs). These siRNAs are loaded into ARGONAUTE 1 (AGO1), forming a complex that interacts with mRNAs by complementarity to the loaded siRNAs. When the siRNA is 21-nt long, targeted RNAs are mostly cleaved and subsequently degraded. However, when the siRNA is 22-nt long, targeted RNAs are transformed into dsRNA by RNA dependent RNA polymerase 6 (RDR6) and SUPPRESSOR OF GENE SILENCING 3 (SGS3) to produce secondary siRNAs, thereby amplifying PTGS. A fraction of the siRNAs can move to neighbouring cells through plasmodesmata or to distant tissues through phloem transport, leading in some cases to virus recovery and transgene systemic PTGS (Palauqui et al. 1997; Voinnet and Baulcombe 1997; Kobayashi and Zambryski 2007; Vogler et al. 2008).

Given that viruses produce high amounts of dsRNA during their replication cycle, it is easy to conceive that PTGS is activated upon infection. However, how sense transgenes that are not supposed to produce dsRNA activate a form of PTGS referred to as sense (S)-PTGS has long been a puzzling question. When S-PTGS of plant host genes induced by homologous sense transgenes was first reported (Napoli et al. 1990; Van Der Krol et al. 1990), emphasis was given to copy number because S-PTGS occurred more often in plants carrying multiple transgene copies. An alternative, although non-exclusive, hypothesis proposed that a precise gene-specific RNA level should be passed to trigger PTGS. This RNA threshold hypothesis was supported by the fact that transgenic lines could be classified in three categories: i) those that express the transgene at very high level and trigger S-PTGS independently of the status of the transgene locus, homozygous or hemizygous, ii) those that express the transgene at lower level and trigger S-PTGS only when homozygous for the transgene locus and not when hemizygous, and iii) those that express the transgene at very low level and never trigger S-PTGS spontaneously (de Carvalho et al. 1992; Hart et al. 1992; de Borne Dorlhac et al. 1994; Palauqui and Vaucheret 1995; Vaucheret et al. 1995). Remarkably, single-copy transgenic lines of the second class triggered PTGS as efficiently in haploid plants as in homozygous diploid plants (Elmayan and Vaucheret 1996), indicating that S-PTGS correlates with the relative quantity of a given RNA per cell and not with the presence of multiple transgene copies. Importantly, the RNA threshold hypothesis was perfectly compatible with the copy number hypothesis, simply because, generally, the higher the number of copies is, the higher is the amount of RNA produced (Schubert et al. 2004).

Despite the fact that the RNA threshold hypothesis is now commonly accepted, it remains unclear if a single RNA threshold conditions all S-PTGS steps or if multiple thresholds corresponding to distinct RNA molecules condition different S-PTGS steps. Transgenic lines in which S-PTGS is initiated locally but does not become systemic have been reported (Kalantidis et al. 2006). Inversely, certain transgenic lines that do not spontaneously trigger S-PTGS can undergo S-PTGS when grafted onto silenced lines, indicating that despite being incompetent for spontaneous S-PTGS initiation, they are totally competent for S-PTGS amplification (Taochy et al. 2017). Taken together, it suggests that S-PTGS initiation and amplification require distinct thresholds.

Supporting this hypothesis, initiation and amplification can be genetically uncoupled. This was exemplified through genetic screens involving the Arabidopsis *Pro35S:GUS* lines *L1* and *L2,* which spontaneously undergoes S-PTGS, as well as the *Pro35S:GUS* line *6b4,* which never triggers S-PTGS spontaneously, but which undergo S-PTGS upon grafting onto line *L1*, indicating that it supports S-PTGS amplification. Although being incompetent to initiate S-PTGS spontaneously in a wildtype background, line *6b4* triggers S-PTGS in mutants impaired in RNA quality control (RQC) components (Gy et al. 2007; Moreno et al. 2013; Martínez de Alba et al. 2015; Yu et al. 2015; Hématy et al. 2016; Lange et al. 2019), suggesting that a certain amount of aberrant RNAs is necessary to initiate S-PTGS. Supporting this hypothesis, analysis of *L1, L2, L1 rdr6, L2 rdr6, 6b4* and *6b4 rdr6* plants revealed the existence of an uncapped RNA antisense to *GUS* referred to as *SUG*. This RNA accumulates in both wildtype and *rdr6* backgrounds, indicating that it is not a downstream product of S-PTGS. Because uncapped RNAs are sensitive to 5’-to-3’ degradation by EXORIBONUCLEASE3 (XRN3) and XRN4, *SUG* accumulation was monitored in *6b4 xrn3 xrn4 rdr6* plants and found higher than in *6b4 rdr6* (Parent et al. 2015). *SUG* accumulation was also higher in *L1 rdr6* and *L2 rdr6* plants compared with *6b4 rdr6.* Therefore, *SUG* was considered a potential aberrant RNA (abRNA) that could serve as substrate for RDR6 to initiate *GUS* S-PTGS when passing a certain threshold or when escaping degradation by RQC, explaining why *L1, L2* and *6b4 xrn3 xrn4* initiate S-PTGS spontaneously while *6b4* does not due to efficient degradation of uncapped *SUG* by XRN3 and XRN4 (Gy et al. 2007).

Another genetic screen identified mutations in the gene encoding the histone H3K4 di/trimethyl demethylase JUMONJI 14 (JMJ14), a protein previously shown to regulate a subset of endogenous genes (Jeong et al. 2009; Yang et al. 2010). Our work showed that the *jmj14* mutation reduces, but does not impair, *L1* S-PTGS. Indeed, *L1 jmj14* plants were still able to initiate S-PTGS and produce a mobile silencing signal (Jeong et al. 2009; Yang et al. 2010; Le Masson et al. 2012; Butel et al. 2021). However, *jmj14* suppressed the capacity of line *6b4* to undergo S-PTGS upon grafting onto line *L1* (Butel et al. 2021), suggesting that *jmj14* affects another S-PTGS step.

Although these data suggested different requirements for S-PTGS initiation and amplification, whether these different requirements corresponded to distinct RNA thresholds has not been determined yet. In this study, we examined separately these requirements, and came to the conclusion that two distinct RNA thresholds exist, which independently condition S-PTGS initiation and amplification.

## Results

### Identification of second site mutations restoring transgene S-PTGS in a jmj14 mutant background

A genetic screen based on the Arabidopsis *Pro35S:GUS* line *L1,* which spontaneously undergoes S-PTGS with 100% efficiency at each generation, identified *jmj14* as a mutation partially suppressing S-PTGS (Le Masson et al. 2012). To further characterize the effect of *jmj14*, the non-silenced Arabidopsis *Pro35S:GUS* line *6b4* was grafted onto *L1* and *L1 jmj14* plants. Similar to *L1, L1 jmj14* rootstocks induced a robust S-PTGS response in grafted *6b4* plants, although slightly delayed, indicating that *jmj14* does not impair S-PTGS initiation or siRNA propagation (Butel et al. 2021). In contrast, *6b4 jmj14* plants did not undergo S-PTGS upon grafting onto line *L1*, suggesting that *jmj14* affects another S-PTGS step (Butel et al. 2021).

To understand how *JMJ14* promotes systemic S-PTGS, a genetic screen was conducted to identify second site mutations restoring S-PTGS in *jmj14*. The *6b4* locus could not be used for such a screen because it would require grafting every mutagenized plant to determine its S-PTGS efficiency. The *L1* locus could be used for such a genetic screen; however, it would be time consuming and would require testing GUS activity in mutagenized plants one by one. To make the screen simpler, the *JAP3* locus was introduced into *L1 jmj14* plants. This locus was chosen because it expresses a hairpin containing part of the *PHYTOENE DESATURASE (PDS)* gene in companion cells (CCs), which causes *PDS* silencing and photobleaching in cells surrounding the veins in a JMJ14-dependent manner (Smith et al. 2007; Searle et al. 2010). We have confirmed that *L1 JAP3 jmj1*4 plants exhibit behaviour similar to *L1 jmj14* and *JAP3 jmj14* plants in terms of *GUS* and *PDS* PTGS (Figures 1A and 1B), which validates the use of this line for conducting the genetic screen.

**Figure 1:**
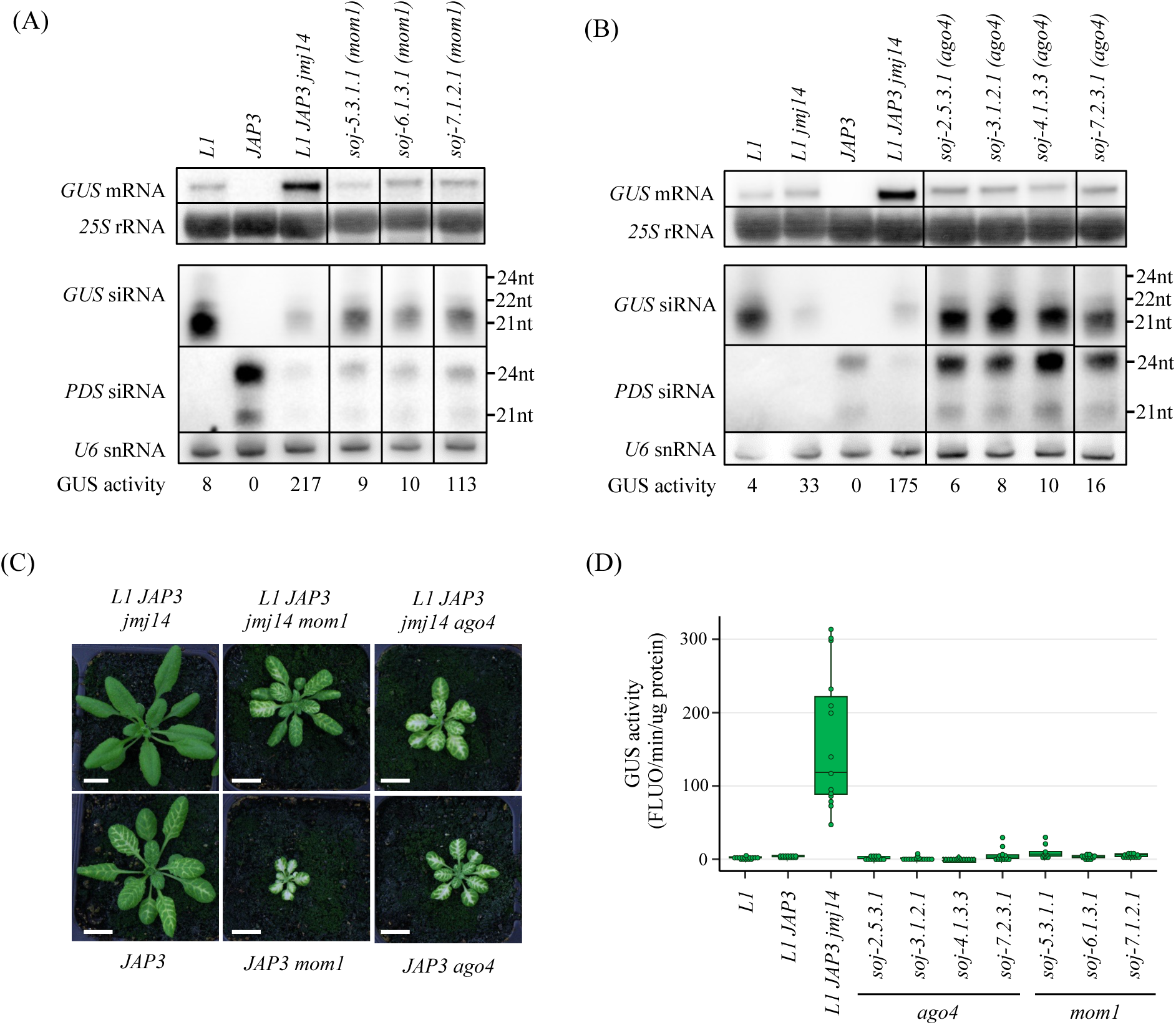
Mutations in *AGO4* and *MOM1* restores *L1* S-PTGS in a *JMJ14*-deficient background. (A) and (B) Northern blot analysis showing *GUS* mRNA and *GUS* and *PDS* siRNA accumulation in 14 days after germination shoots of the indicated genotypes. *25S* rRNA and *U6* snRNA probes were used for normalization. GUS activity is in arbitrary units of fluorescence. µg of protein^−1^.min^−1^. Uncropped blots can be seen in the Supplemental Figure 2. (C) Phenotype of plants carrying the *ProSUC2:hpPDS* locus *JAP3* in the indicated genotypes. Scale bar, 1cm. (D) GUS activity in leaves of plants carrying the *Pro35S:GUS* locus *L1* in the indicated genotypes. Mutants recovered from the mutagenesis of *L1 JAP3 jmj14* plants were originally referred to as *soj (suppressor of jmj14).* Each dot represents an individual plant, with more than 10 plants tested per experiment.

*L1 JAP3 jmj14* seeds were EMS mutagenized, and 16 M2 plants exhibiting restored photobleaching were identified (Figure 1C, Supplemental Figures 1A). Among these 16 M2 plants, seven showed increased levels of both *PDS* siRNA and *GUS* siRNA compared to *L1 JAP3 jmj14*, as well as reduced levels of *GUS* mRNA and GUS activity (Figures 1). Efficient restoration of *GUS* PTGS was confirmed in these seven mutants in the M3 generation (Figure 1D).

### Mutations in AGO4 and MOM1 restore S-PTGS in a JMJ14-deficient background

Because *ago4* was previously shown to suppress the effect of *jmj14* on *PDS* silencing mediated by the *JAP3* locus (Searle et al. 2010), and *dcl3* to enhance *PDS* silencing mediated by the *JAP3* locus (Smith et al. 2007), complementation tests were performed on the seven mutants showing restored *GUS* S-PTGS to determine if they correspond to *ago4* or *dcl3* alleles (Supplemental Figures 1B-C). Four mutants of the seven mutants corresponded to *ago4* alleles. To determine which mutations were present in the three remaining mutants, whole-genome sequencing was performed, revealing that they correspond to *mom1* alleles. The position and the effect of the identified mutations are indicated in the Supplemental Figure 1D.

To determine if *ago4* and *mom1* mutations are causal in the restoration of systemic S-PTGS in a *jmj14* mutant background, *6b4 ago4, 6b4 jmj14 ago4*, *6b4 mom1* and *6b4 jmj14 mom1* plants were generated. Like *6b4* and *6b4 jmj14* controls, none of these plants triggered S-PTGS spontaneously, indicating that *ago4* and *mom1* do not promote S-PTGS initiation (Figure 2A). In contrast, systemic S-PTGS was observed in *6b4, 6b4 ago4, 6b4 jmj14 ago4*, *6b4 mom1* and *6b4 jmj14 mom1* plants grafted onto *L1* (Figure 2B), indicating that *ago4* and *mom1* are actually capable of restoring systemic S-PTGS in a *jmj14* mutant background, at least at the *L1* and *6b4* loci.

**Figure 2:**
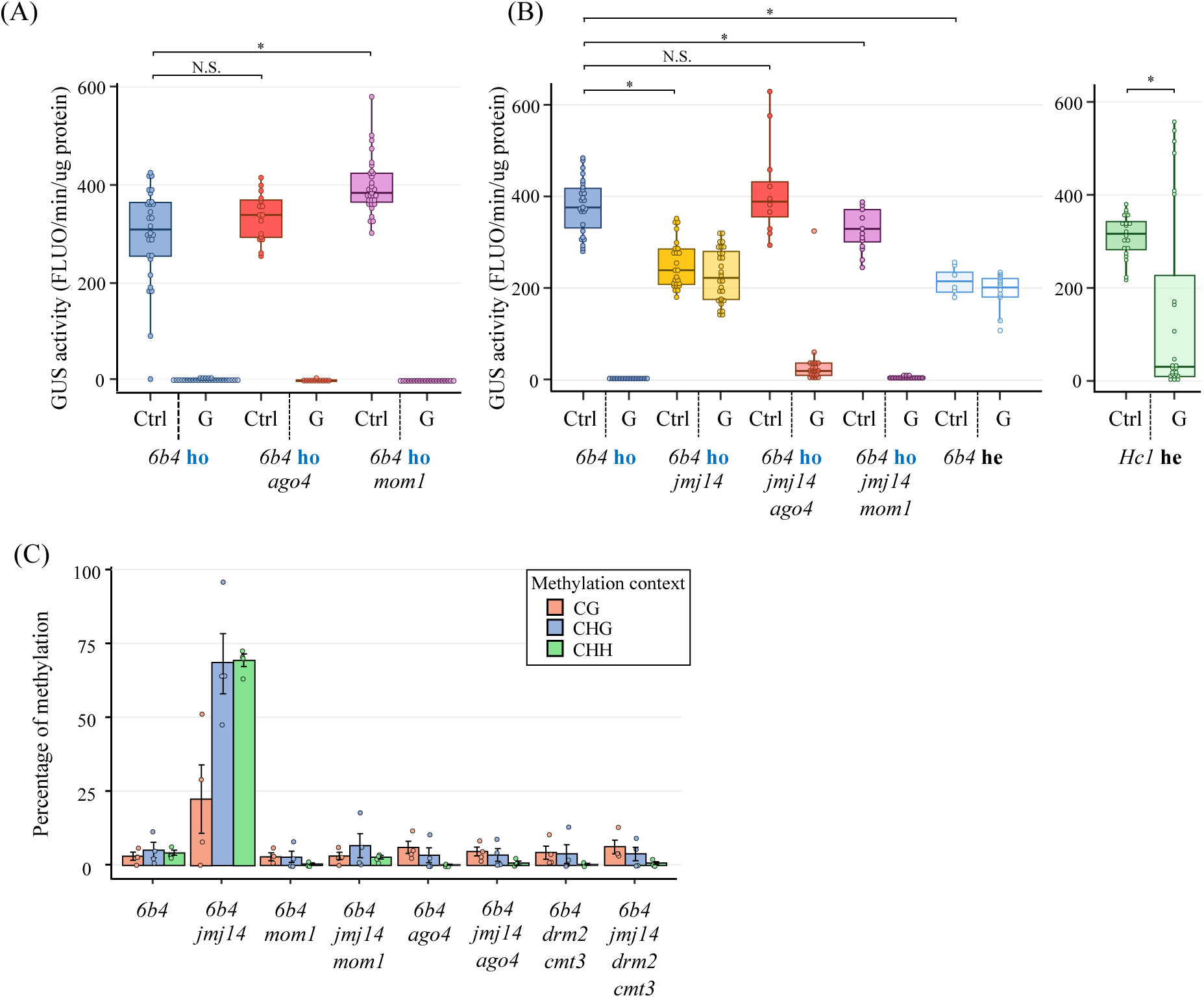
Mutations in *AGO4* and *MOM1* suppress ectopic DNA methylation triggered by *jmj14* at the *6b4* locus and restore its ability to undergo S-PTGS upon grafting onto the silenced *L1* line, similar to mutations in *CMT3* and *DRM2*. (A) and (B) GUS activity in aerial parts of 14 days after germination (dag) plants of the indicated genotypes, either grown on their own roots (Ctrl) or grafted (G) onto an *L1* rootstock. Each dot represents an individual plant, with more than 12 plants tested per experiment. Allelic state is indicated as following: **ho** = homozygous, **he** = hemizygous. For (B), *Hc1* heterozygous grafting experiment has been made independently of the other graft experiments and thus shown apart. P-values were calculated using non parametric Wilcoxon tests adjusted with Holm–Bonferroni correction, except for *Hc1* where non parametric Mann-Whitney test was used (* means P<0.05 and N.S., not significant). (C) DNA methylation analysis of the *35S* promoter at the *6b4* locus in the three different methylation context (CG, CHG and CHH) assessed by Chop-qPCR. Dots represent biological replicates, composed of 4 independent pools of plants.

### Mutations in AGO4 and MOM1 suppress ectopic methylation caused by JMJ14 deficiency

Previous whole-genome bisulfite analysis revealed that the *jmj14* mutation has no impact on the DNA methylation status of endogenous genes, but slightly increases the level of DNA methylation at CG sites in transposable elements (TE) (Butel et al. 2021). In contrast, a striking increase in DNA methylation at CHG and CHH sites was observed at the transgene loci *6b4* and *L1* in a *jmj14* mutant background (Le Masson et al. 2012; Butel et al. 2021), and to a lesser extend at CG sites, similar to what is observed at TE loci. This transgene DNA hypermethylation caused by the absence of JMJ14 was proposed to prevent systemic S-PTGS at transgene loci because *6b4 jmj14 drm2 cmt3* plants exhibited levels of DNA methylation similar to *6b4* and triggered systemic S-PTGS as efficiently as *6b4* plants upon grafting onto *L1* rootstocks, indicating that JMJ14 is not required for systemic S-PTGS (Butel et al. 2021).

The importance of AGO4 and MOM1 for de novo DNA methylation was previously revealed when newly introduced *FWA* transgenes failed to become methylated in RdDM mutants, including *ago4* (Chan et al. 2004), and in the *mom1* mutant (Li et al. 2023). The identification of *ago4* and *mom1* as second-site suppressors of *jmj14* therefore suggested that, similar to *drm2 cmt3, ago4* and *mom1* could suppress transgene ectopic DNA methylation induced by *jmj14*, allowing *6b4 jmj14 ago4* and *6b4 jmj14 mom1* plants to undergo S-PTGS upon grafting because transgene DNA methylation is back to *6b4* levels. Quantification of DNA methylation levels by methylation-sensitive DNA digestion followed by qPCR shows that the ectopic DNA methylation of the *35S* promoter observed in *6b4 jmj14* was actually lost in *6b4 jmj14 ago4* and *6b4 jmj14 mom1*, similar to what is observed in *6b4 jmj14 drm2 cmt3* (Figure 2C). These results therefore confirm a correlation between the ability of the *Pro35S:GUS* to undergo *S-PTGS* and its DNA methylation status.

### Transgene S-PTGS is indirectly suppressed by DNA hypermethylation

The fact that ectopic transgene DNA methylation is observed in *6b4 jmj14* plants, which are deficient for S-PTGS, and not in *6b4 jmj14 drm2 cmt3*, *6b4 jmj14 ago4* and *6b4 jmj14 mom1* plants, which are capable of triggering S-PTGS upon grafting, does not mean that DNA methylation plays a direct role. Remarkably, GUS activity in *6b4 jmj14* is lower than in *6b4*, whereas it is similar to *6b4* in *6b4 jmj14 ago4* and *6b4 jmj14 mom1* (Supplemental Figure 3). Therefore, the incapacity of *6b4 jmj14* to trigger systemic S-PTGS could result either directly from transgene DNA hypermethylation or indirectly from a reduced level of transgene mRNA due to DNA hypermethylation induced in the *35S* promoter when JMJ14 is absent. To discriminate between these two hypotheses, plants heterozygous for the *6b4* locus (*6b4*he) were generated by crossing plants homozygous for *6b4* (*6b4*ho) to wildtype plants. *6b4*he plants exhibited two-fold less GUS activity compared to *6b4*ho (Figure 2B), and were at a level similar to *6b4 jmj14*. Upon grafting onto *L1* rootstocks, *6b4*he plants did not trigger S-PTGS (Figure 2B), strongly suggesting that *6b4 jmj14* plants do not trigger systemic S-PTGS upon grafting because of an insufficient level of target mRNAs. Together, these results strongly suggest that, in *jmj14*, systemic S-PTGS is impaired owing to the effect of transgene promoter DNA hypermethylation on the transcription of transgene mRNA.

### Analysis of a very large set of transgene loci confirms the correlation between transgene expression and systemic S-PTGS

The set of *Pro35S:GUS* transgenic lines originally characterized (Elmayan et al. 1998) comprised line *6b4*, which does not trigger S-PTGS spontaneously, lines *L1* and *L2*, which trigger S-PTGS spontaneously with 100% efficiency independent of the homozygous (ho) or hemizygous (he) status of the transgene, and line *Hc1*, which triggers S-PTGS in 20% of the homozygous plants at each generation but not in hemizygous plants. To further precise the transcriptional threshold level that conditions the spontaneous triggering of S-PTGS, GUS activity was quantified as a proxy for *GUS* mRNA levels, in *L1*ho, *L2*ho, *Hc1*ho, *Hc1*he, *6b4*ho and *6b4*he plants in wildtype or PTGS-deficient (*rdr6* or *sgs3*) backgrounds. A perfect correlation was observed between the level of GUS activity in non-silenced plants and the efficiency of systemic S-PTGS (Figures 3A). Indeed, *L1*ho plants, which trigger S-PTGS with 100% efficiency and the most rapidly during development, show the highest level of GUS activity, defined in a *rdr6* mutant background (Figure 3A). Line *L2*ho, which also triggers S-PTGS with 100% efficiency but less rapidly than line *L1 (*Figure 3A), shows the second highest level of GUS activity. Line *Hc1*, which triggers S-PTGS in 20% of the population, exhibits GUS activity in the non-silenced fraction of the population or in the *sgs3* background below that in *L2*ho *rdr6* and *L1*ho *rdr6* (Figure 3A), but higher than in lines *6b4*ho, *Hc1*he and *6b4*he plants, which never trigger S-PTGS spontaneously. Among this last category, *6b4*ho plants trigger S-PTGS with 100% efficiency upon grafting onto *L1*, *Hc1*he plants with 50% efficiency, whereas *6b4*he plants do not trigger S-PTGS at all (Figure 2B), correlating perfectly with their expression level (Figure 3A). Together, these results indicate that the threshold for systemic S-PTGS is just above the level of transgene expression in *6b4*he and close to the level in *Hc1*he plants.

**Figure 3:**
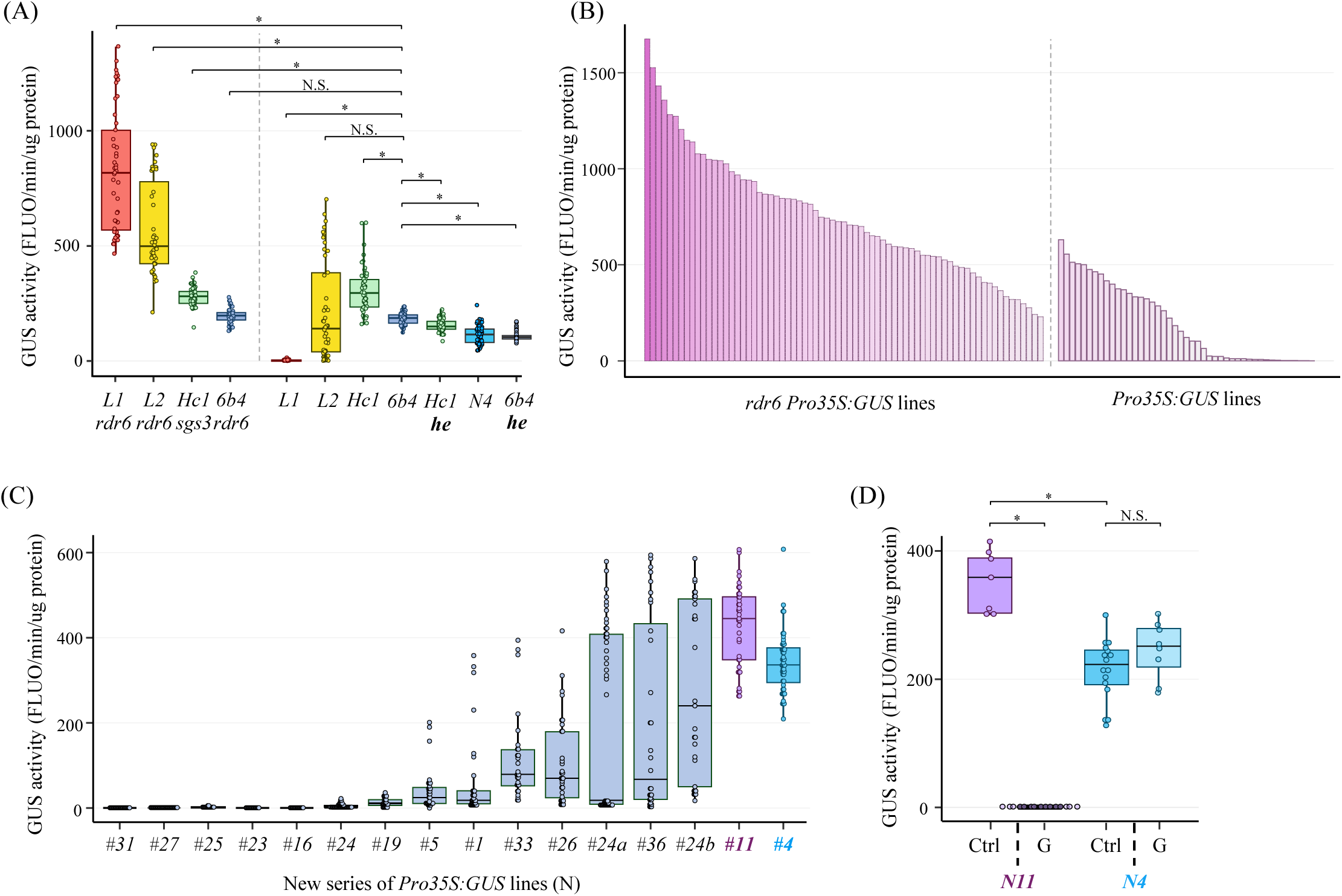
Analysis of a large series of *Pro35S:GUS* lines reveals a discrete amplification threshold. (A) GUS activity of *Pro35S:GUS-tRbcS/tNOS-NPTII:pNOS* (*Pro35S:GUS*) lines in WT or PTGS-deficient mutant background (*rdr6* or *sgs3*). GUS activity has been measured in aerial parts of plants of 11 days after germination in long-day conditions. Each dot represents an individual plant. Three biological replicates were tested for a total of 36 plants per line. P-values were calculated using non parametric Wilcoxon tests adjusted with Holm–Bonferroni correction (* means P<0.05 and N.S., not significant). (B) GUS activity of a new series of primary transformants obtained by introducing the *Pro35S:GUS* construct in either WT Col-0 or *rdr6* mutant. Each bar represents an individual transformant. (C) GUS activity in homozygous T3 plants of several WT / *Pro35S:GUS* lines of the new series carrying the T-DNA inserted at a single locus. Each dot represents an individual, with at least 16 plants tested per line. (D) GUS activity in homozygous plants of *line N11* and *line N4* grafted (G) or not (Ctrl) onto *L1* roots. P-values were calculated using non parametric Wilcoxon tests adjusted with Holm–Bonferroni correction (* means P<0.05 and N.S., not significant).

To extend this analysis, we generated a large number of new transformants by introducing the *Pro35S:GUS* construct into Col-0 or *rdr6*. Analysis of T1 transformants revealed that all *rdr6*/*Pro35S:GUS* transformants (n = 73) expressed GUS (Figure 3B), whereas only 50% of Col-0/*Pro35S:GUS* transformants (n = 47) exhibited significant GUS activity, i.e. above that observed in silenced *L1* plants. Remarkably, the maximum GUS activity observed in Col-0/*Pro35S:GUS* transformants corresponded to a level close to that in *6b4*ho plants (although plants were not tested at the exact same stage of development), strongly suggesting that when expressed above a certain level, transgene loci spontaneously trigger S-PTGS. To examine further the relationship between transgene expression and S-PTGS efficiency, non-silenced Col-0/*Pro35S:GUS* T1 transformants carrying the T-DNA inserted at a single locus, i.e. showing a 3:1 segregation ratio, were brought to the homozygous T3 stage. Because homozygous T3 plants express the transgene at twice the level of the T1 transformants, a large proportion of the unsilenced T1 lines were expected to trigger PTGS in a homozygous state, at least in a fraction of the population. Analysis of GUS activity in the 16 lines carrying the T-DNA inserted at a single locus (Figure 3C) revealed that 7 lines trigger silencing in 100% of the population (similar to the original lines *L1* and *L2*), 7 lines trigger silencing in a variable fraction of the population (similar to the original line *Hc1*), whereas 2 lines do not trigger S-PTGS spontaneously (similar to the original line *6b4*). Among these last two lines, one (line *N11*) expressed GUS at a level close to that of *6b4*ho plants and, like *6b4*ho, triggered S-PTGS with 100% efficiency upon grafting onto *L1*, whereas the second line (line *N4*) expressed GUS at a level close to that of *6b4*he plants (Figure 3A) and, like *6b4*he, did not trigger PTGS upon grafting onto *L1* (Figure 3D).

Overall, the analysis of this very large set of additional transgenic lines indicates that the vast majority of *Pro35S:GUS* loci are able to trigger S-PTGS spontaneously (at frequencies ranging from 1% to 100%), and that only a very limited number of lines are unable to initiate S-PTGS spontaneously in a wild-type background. Among these, only one line (*N4*) appears incapable to undergo systemic S-PTGS upon grafting onto *L1*, while the other (*N11*) behaves like *6b4* and undergoes S-PTGS upon grafting onto *L1*. Importantly, the behaviour of the line *N11* indicates that the uncoupling of S-PTGS spontaneous initiation and graft-induced amplification is not a specific trait of line *6b4*. Rather, these results suggest the existence of distinct thresholds conditioning S-PTGS initiation and amplification. Most *Pro35S:GUS* lines are above these two thresholds, whereas lines *6b4* and *N11* are above the S-PTGS amplification threshold but below the S-PTGS initiation threshold, and line *N4* below both thresholds.

### Transient passing of the S-PTGS initiation threshold allows triggering systemic S-PTGS

Previous analysis of S-PTGS using a visual system based on the cosuppression of endogenous *NIA* (*NITRATE REDUCTASE*) or *NII* (*NITRITE REDUCTASE*) genes by *Pro35S*-driven transgenes revealed that spontaneous S-PTGS is always visualized as a single spot on a single leaf before spreading throughout the plant (Palauqui et al. 1996), suggesting that a single and likely transient event of initiation is sufficient to trigger systemic S-PTGS. To test this hypothesis, we designed a system allowing an additional *Pro35S:GUS* transgene copy to be transiently expressed on top of the stably expressed *Pro35S:GUS* locus *6b4*. For this purpose, an estradiol-inducible Lex/*Pro35S:GUS* transgene (referred to as *iGUS*) was generated and introduced into line *6b4*. It is well known that estradiol-inducible constructs can sometimes leak, but this generally has no consequence if it happens accidentally in a few cells. However, in the case of a transgenic line supporting S-PTGS amplification, leakage in a few cells could translate into systemic S-PTGS. 50% of the transformants triggered *GUS* PTGS (n=9/18 plants) when grown in absence of estradiol, suggesting that most *iGUS* loci leak, at least in some cells, in the absence estradiol, and that the increase of *GUS* expression occurring in these leaking cells triggers systemic *GUS* S-PTGS. Another issue with estradiol-inducible constructs is that lines that do not leak at all hardly express the construct upon induction. Among the T1 plants that did not trigger *GUS* S-PTGS in the absence of estradiol, only one showed no triggering of *GUS* S-PTGS in T2 plants grown in the absence of estradiol, but efficient triggering of *GUS* S-PTGS on media supplemented with estradiol (Figure 4A). We took advantage of the fact that estradiol is unstable and that estradiol induction does not last after transferring plants to estradiol-free medium (Zuo et al. 2000) to determine if a transient induction is sufficient to induce S-PTGS. For this purpose, homozygous T3 plants were sown on media supplemented with estradiol and transferred to estradiol-free media after 0, 3, 9, 15 or 21 days. *GUS* PTGS quantification revealed that 3 days on estradiol was sufficient to trigger *GUS* PTGS, but that the longer plants were left on estradiol, the higher was the frequency of plants triggering *GUS* PTGS (Figure 4B), confirming that increasing *GUS* RNA level in the *6b4* line allows passing the *GUS* PTGS initiation threshold. The exact increase of GUS activity was determined by two independent ways. One consisted in segregating the *6b4* locus and growing plants containing only the *iGUS* locus on media with or without estradiol. The second consisted in crossing the *6b4/iGUS* line with *6b4 rdr6* plants and selecting F2 homozygous for the three loci (*6b4, iGUS, rdr6*), and comparing GUS activity in *6b4/iGUS rdr6* vs *6b4 rdr6* plants grown with or without estradiol. Both ways showed that the *iGUS* locus contributed approximately one fourth of the activity of the *6b4* locus (Figure 4C), indicating that a slight increase above the *6b4* level is sufficient to pass the initiation threshold. The results obtained with the *6b4/iGUS* plants treated transiently with estradiol also indicate that once initiated by a pulse of transgene over-expression, systemic S-PTGS can be maintained in the absence of the inducing agent. This was suggested but could not be experimentally tested by grafting experiments because it required separating the scions from the silenced rootstock after PTGS induction.

**Figure 4:**
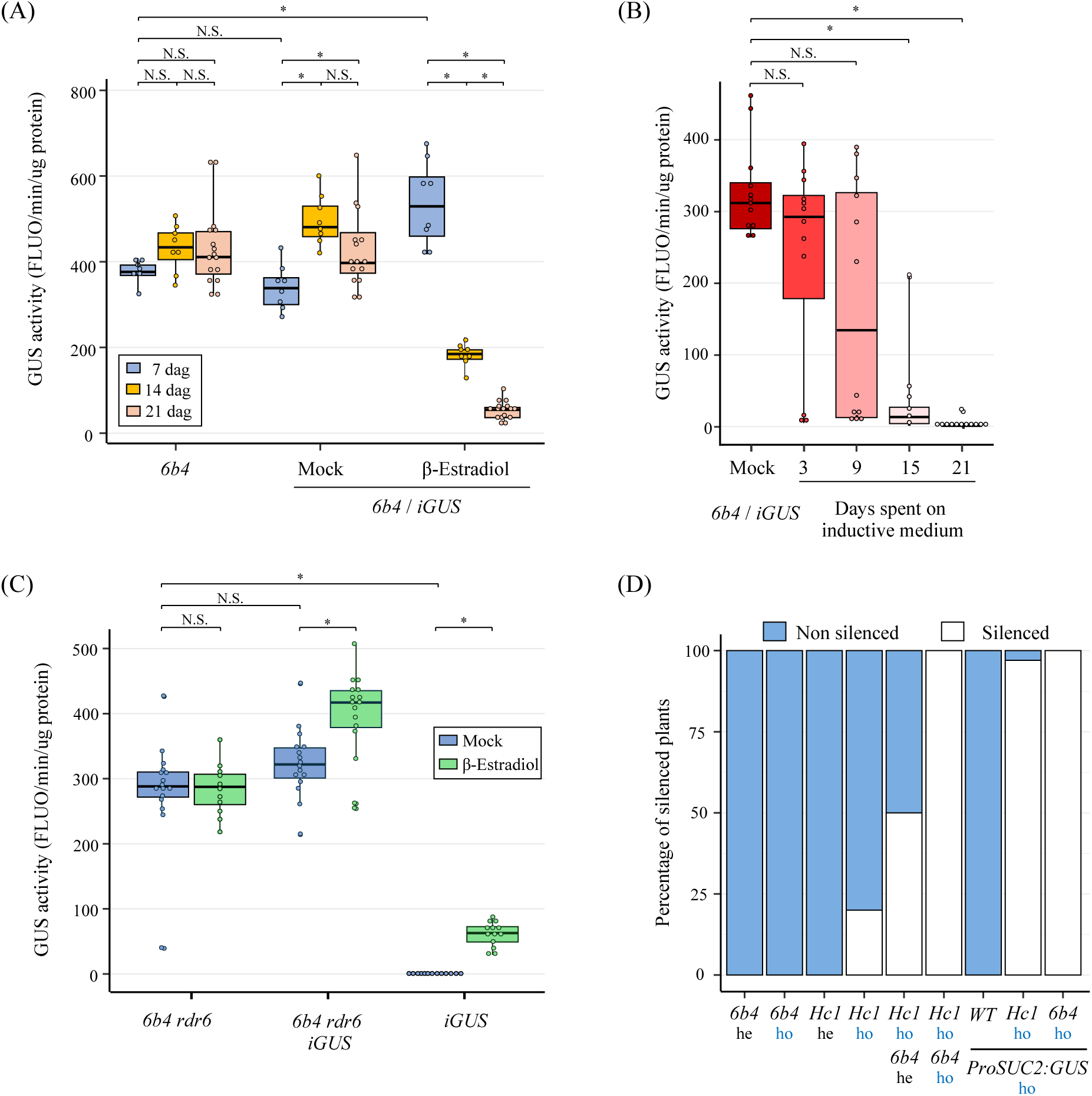
Local or transient increase of GUS expression in the *6b4* line triggers systemic S-PTGS. (A) GUS activity in leaves of *6b4* controls and T3 homozygous plants of a *6b4/iGUS* line obtained by introduction of an inducible *GUS* construct in the *6b4* line. Seeds were sown on medium supplemented or not with 20µM β-estradiol and sampled at 7, 14 and 21 days after germination (dag). Each dot represents an individual plant, and more than eight plants were tested per condition. P-values were calculated using a two way Scheirer-Ray-Hare test adjusted with Holm–Bonferroni correction (* means P<0.05 and N.S., not significant). (B) T3 homozygous *6b4*/*iGUS* seeds were sown on media supplemented with 20µM β-estradiol before being transferred to media without β-estradiol at 0, 3, 9, 15 or 21 dag. At 21dag, plants were transferred to soil. GUS activity was quantified from leaves 3 weeks later. Each dot represents an individual plant, and more than 12 plants were tested per condition. P-values were calculated using non parametric Wilcoxon tests adjusted with Holm–Bonferroni correction (* means P<0.05 and N.S., not significant). (C) GUS activity in leaves of the *6b4/iGUS* line in a *rdr6* background compared to *6b4 rdr6,* and in the *iGUS* line from which the *6b4* locus has been segregated. Lines were treated with 20µM β-estradiol or not (mock) to assess the level of GUS activity produced by the *iGUS* locus. Each dot represents an individual plant, and more than 13 plants were tested per condition. P-values were calculated using a two way Scheirer-Ray-Hare test adjusted with Holm–Bonferroni correction (* means P<0.05 and N.S., not significant). (D) Proportion of *GUS* silenced (S) and non-silenced (NS) plants in the *Pro35S:GUS* lines *6b4 and Hc1* in different allelic states (**ho** = homozygous, **he** = hemizygous) either alone or combined with a *ProSUC2:GUS* line expressing *GUS* in phloem companion cells. Plants were considered silenced when *GUS* expression was less than 5% of non-silenced controls. 96 plants were analyzed per genotype.

### Transgenes driven by non-viral promoters also allow passing the S-PTGS initiation threshold

With the exception of one transgenic plant carrying an ectopic copy of the endogenous petunia *CHS* gene (Van Der Krol et al. 1990), all known S-PTGS events result from the introduction of transgenes driven by the viral promoter *Pro35S*. Here we showed that expressing a transgene based on the human/plant virus chimeric promoter *Lex*/*Pro35S* also contributes to reaching the amount of abRNAs that triggers S-PTGS. To determine if a foreign origin of the promoter is key to initiating S-PTGS, the effect of a *GUS* transgene driven by a plant promoter was analyzed. For this purpose, a *ProSUC2:GUS* locus expressing *GUS* only in the phloem companion cells (Brioudes et al. 2021) was combined with different *Pro35S:GUS* loci. To ensure that this *ProSUC2:GUS* line does not spontaneously produce mobile siRNAs, the *6b4* line was grafted onto the *ProSUC2:GUS* line. None of the grafted plants triggered S-PTGS, confirming that the *ProSUC2:GUS* line does not spontaneously produce siRNAs that could move to line *6b4* and induce S-PTGS amplification (Supplemental Figure 4). Then, line *ProSUC2:GU*S was crossed to lines *Hc1* and *6b4*. Plants homozygous for *ProSUC2:GUS* and *Hc1* or *ProSUC2:GUS* and *6b4* were identified in the F2 progeny, and F3 populations were analyzed. 100% of the *Hc1 ProSUC2:GUS* (n = 96) and 97% of the *6b4 ProSUC2:GUS* plants (n = 96) triggered S-PTGS (Figure 4D). Given that neither *6b4* nor *ProSUC2:GUS* triggers S-PTGS spontaneously, these results indicate that combining *6b4* with *ProSUC2:GUS* allows reaching a level of aberrant RNAs sufficient to pass the S-PTGS initiation threshold, similar to combining *6b4* with the PTGS-prone locus *Hc1* (Figure 4D). Moreover, the fact that *ProSUC2:GUS*, which is expressed in only one cell type, the phloem companion cells, promotes *6b4* S-PTGS and enhances *Hc1* S-PTGS confirms that a localized initiation by the combined loci is sufficient to trigger systemic S-PTGS.

### Impairing DCL4 promotes systemic S-PTGS on endogenous genes

The results presented in this study support the hypothesis of the existence of distinct thresholds conditioning S-PTGS initiation and amplification. Likely, the former is related to the amount of abRNAs that escape RQC to be transformed into siRNAs, while the latter is related to the amounts of mRNAs targeted by siRNAs. However, siRNAs involved in S-PTGS come in two flavors. One class is 21-nt long and is produced by DCL4 while the other is 22-nt long and is produced by DCL2. 21-nt siRNAs only guide the cleavage of target mRNA while 22-nt siRNAs engage these cleaved mRNAs in the production of additional dsRNAs by RDR6 and subsequent amplification of siRNAs, producing the so-called secondary siRNAs. Long ago, the balance between DCL2 and DCL4 was shown to condition the efficiency of transgene S-PTGS, *dcl2* mutants showing decreased S-PTGS and *dcl4* mutants increased S-PTGS (Mlotshwa et al. 2008; Parent et al. 2015; Taochy et al. 2019). This is due to the fact that, in wildtype plants, DCL2-dependent 22-nt siRNAs represent a minority compared to DCL4-dependent 21-nt siRNAs because DCL4 somehow obscures the action of DCL2.

Like transgenes, endogenous genes produce abRNAs (Martinez de Alba et al., 2015; Zhang et al., 2015). However, siRNAs originating from endogenous genes are only detected in RQC-deficient mutants because evolution has shaped RQC to allow an efficient degradation of endogenous abRNAs to avoid the eventual silencing of endogenous genes. Nevertheless, it is reasonable to assume that unintended RQC dysfunction can occur locally after a stress or for other reasons, and that endogenous genes produce siRNAs in some cells. Given the high level of expression of certain endogenous genes, these locally-produced siRNAs could potentially trigger systemic S-PTGS. To determine if endogenous genes have the potential to undergo systemic S-PTGS after local initiation and if DCL4 prevents this to happen, WT and *dcl4* scions were grafted onto transgenic rootstocks expressing a *Pro35S:NIA2* transgene (*2a3*), which causes S-PTGS of the endogenous *NIA1* and *NIA2* genes (Figure 5A-B). These endogenous genes were chosen because i) they are highly expressed, and ii) they are in the top 10 of the endogenous genes producing siRNAs in RQC-deficient mutants (Martínez de Alba et al. 2015; Zhang et al. 2015). Moreover, the production of *NIA1* and *NIA2* siRNAs is enhanced in double mutants between *dcl4* and RQC-deficient mutants (Feng et al. 2024). Because *Pro35S:NIA2* lines die at an early stage due to their incapacity to assimilate nitrate, a *Pro35S:NIA2* line in a *jmj14* background (*2a3 jmj14*) was used as rootstock (Le Masson et al. 2012). This line still produces *NIA* siRNAs, which are almost exclusively 21-nt in length, indicating that they are preferentially produced by DCL4 (Figure 5C). Nevertheless, the *2a3 jmj14* line grows sufficiently to allow grafting. None of the WT plants grafted onto *2a3 jmj14* exhibited systemic *NIA* S-PTGS, confirming that despite their capacity to produce siRNAs when RQC is impaired, *NIA1* and *NIA2* genes are incapable of amplifying S-PTGS in a systemic manner in wildtype plants. In contrast, despite being totally free of *NIA* transgenes, *dcl4* mutants efficiently triggered systemic *NIA* PTGS when grafted onto *2a3 jmj14* rootstocks, and accumulated *NIA* siRNAs at high level, which were almost exclusively 22-nt in length, as expected when the absence of DCL4 allows DCL2 to process RDR6 products (Figure 5C). Graft-induced systemic S-PTGS was reproduced using three different *dcl4* alleles (Supplemental Figure 5). These results indicate that, like transgenes, highly expressed endogenous genes such as the *NIA* genes are prone to undergo systemic S-PTGS. However, the fact that *dcl4* mutants never trigger systemic *NIA* S-PTGS spontaneously indicates that RQC is highly efficient at preventing S-PTGS initiation by destroying *NIA* abRNAs, even if DCL4 is non-functional. Reciprocally, the natural DCL2/DCL4 balance prevents S-PTGS to become systemic if RQC accidentally becomes non-functional in certain cells of the plant.

**Figure 5:**
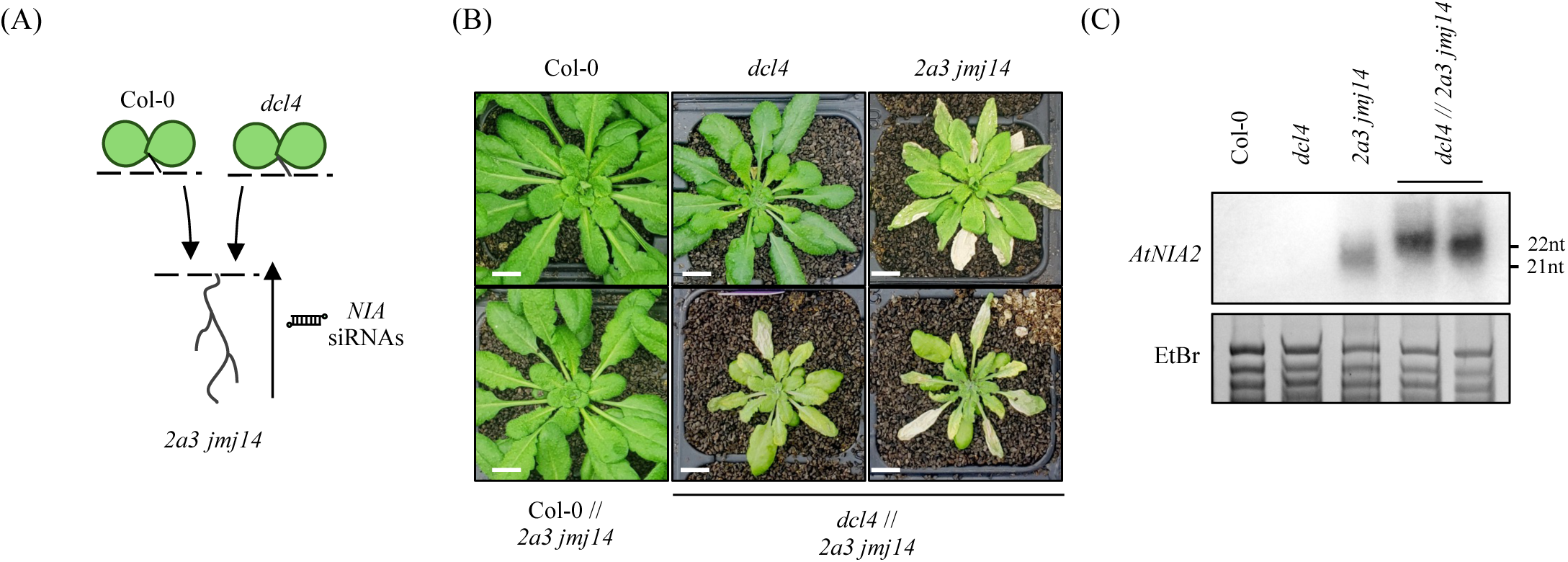
Mutations in *DCL4* allow endogenous *NIAs* genes to trigger systemic PTGS upon grafting. (A) Col-0 or *dcl4-5* mutant scions were grafted onto a silenced *Pro35S:NIA2* transgenic line (*2a3*) rootstock used as an *NIA* siRNAs source. Since plants carrying the *2a3* locus in a Col-0 background die, *2a3 jmj14* plants were used as rootstocks. As *jmj14* reduces NIA silencing without completely impairing *NIA* siRNA production, it allows the rootstocks to survive. (B) Photos showing Col-0 and *dcl4-5* aerial plants grown on their own roots or grafted onto *2a3 jmj14* rootstocks. *2a3 jmj14* plants are shown as PTGS-positive controls. Scale bars, 1cm. (C) Low molecular weight RNA blot analysis of the accumulation of *NIA* siRNAs in *2a3 jmj14* and *dcl4-5* // *2a3 jmj14* leaves aerial parts. Ethidium bromide (EtBr) is used as loading control.

## Discussion

The genetic screen performed on the silenced *Pro35S:GUS* line *L1* retrieved mutations in the core components of the S-PTGS pathway (*AGO1*, *HEN1*, *RDR6*, *SDE5*, *SGS3*) with the exception of *DCL2* and *DCL4* due to their partially redundant action. Unexpected mutants were also retrieved, in particular *jmj14*, whose role first remained elusive. Our whole-genome bisulfite analysis revealed that, contrasting the DNA methylation-independent repressive role of JMJ14 on a subset of endogenous genes, JMJ14 prevents ectopic DNA methylation at transgene loci, thus promoting transgene transcription and S-PTGS (Le Masson et al. 2012; Butel et al. 2021). Moreover, our previous work showed that JMJ14 is not required for the initiation or the execution of PTGS, as siRNAs are still produced in the mutant and can still be transported throughout the plant to mediate a S-PTGS response (Butel et al. 2021). Indeed, scions of the non-silenced *Pro35S:GUS* line *6b4* grafted onto *L1* j*mj14* rootstocks trigger S-PTGS like *6b4* scions grafted onto *L1* (Butel et al. 2021). In contrast, *6b4 jmj14* scions grafted onto *L1* rootstocks do not trigger S-PTGS, indicating that JMJ14 is not absolutely necessary for S-PTGS of lines like *L1*, which spontaneously undergo systemic S-PTGS, but is required for S-PTGS of lines like *6b4*, which do not spontaneously undergo systemic S-PTGS (Butel et al. 2021). Our previous studies also revealed that JMJ14 positively regulates the transcription of *Pro35S:GUS* transgenes at both *L1* and *6b4* loci. JMJ14 binds to the *35S* promoter and recruits NAC52, which somehow inhibits DNA methylation, thereby promoting transcription (Le Masson et al. 2012; Butel et al. 2017, 2021). Suppressing *jmj14*-induced ectopic DNA methylation of the *35S* promoter by the *drm2* and *cmt3* mutations is sufficient to restore S-PTGS in *6b4 jmj14 drm2 cmt3* scions grafted onto *L1* rootstocks, indicating a direct link between DNA methylation, transcription, and systemic S-PTGS.

To further characterize the genetic pathway regulating systemic PTGS, we conducted a suppressor screen on the *L1 JAP3 jmj14* transgenic line, the *JAP3* locus being used as a proxy for easy screening of mutations suppressing the effect of *jmj14* (Figure 1). This screen identified mutations in *MOM1* and *AGO4,* two genes that have been associated with the RNA-directed DNA methylation (RdDM) pathway for repressing the transcription of endogenous RdDM targets as well as inducing TGS on ectopically introduced RdDM-prone transgenes such as *FWA* (Zilberman et al. 2003; Chan et al. 2004; Qi et al. 2006; Yokthongwattana et al. 2010; Li et al. 2023). Not only *L1 JAP3 jmj14 ago4* and *L1 JAP3 jmj14 mom1* showed restored *GUS* and *PDS* PTGS, but *6b4 jmj14 ago4* and *6b4 jmj14 mom1* grafted onto *L1* also showed systemic PTGS similar to *6b4* grafted onto *L1* (Figure 2). Notably, *6b4 ago4* and *6b4 mom1* plants did not trigger S-PTGS spontaneously, indicating that these mutations do not enhance S-PTGS initiation capacities, but enhance S-PTGS amplification capacities, likely by counter-balancing the effect of *jmj14*. Similar to *drm2 cmt3* mutants, *ago4* and *mom1* mutants suppressed the ectopic DNA methylation induced on transgenes by *jmj14*, indicating that AGO4 and MOM1 participate to establishing de novo DNA methylation, irrespective of its mode of induction. The suppression of transgene DNA methylation in *jmj14 ago4* and *jmj14 mom1* coincided with a restoration of transgene transcription at wildtype level, reinforcing the existence of a link between DNA methylation, transcription and S-PTGS amplification (Figure 2). To determine whether DNA methylation or transcription influences S-PTGS amplification, we used heterozygous *6b4* plants, which express *GUS* at levels relatively similar to those in *6b4 jmj14* plants, but that do not carry any mutations or ectopic DNA methylation. Similar to *6b4 jmj14*, heterozygous *6b4* plants failed to trigger systemic S-PTGS when grafted onto *L1* roots (Figure 2). This result implies that S-PTGS is conditioned by the transcription rate, rather than by the DNA methylation status, although the second can influence the first.

The analysis of a large number of independent transformants confirmed the direct relationship between *GUS* level and systemic PTGS as lines exceeding a threshold value triggered a S-PTGS response. They also reveal how rare are *Pro35S:GUS* loci unable to undergo systemic S-PTGS upon grafting onto a silenced rootstock (Figure 3). Together, these results indicate that the level of transgene transcription is critical, and reveal the existence of a discrete mRNA threshold conditioning systemic PTGS.

The fact that line *6b4* is incapable to initiate S-PTGS but is prone to undergo systemic S-PTGS upon grafting onto a silenced rootstock calls for the existence of distinct thresholds conditioning S-PTGS initiation and amplification. Whereas the former is likely related to the amount of aberrant RNAs produced by the locus, the latter is related to the amount of target mRNAs. The probability that a transgene initiates S-PTGS is likely determined by two factors: the propensity of a given locus to produce aberrant RNAs, which depends on its genomic environment but also on the arrangement of the integrated DNA, and its transcription rate, which also depends on its genomic environment and on the number of integrated copies. Line *6b4* produces sufficient *GUS* mRNAs to sustain systemic S-PTGS but not enough abRNAs to initiate S-PTGS. Combining the *6b4* locus with various *GUS* loci expressed in a tissue-specific or inducible manner allows passing the PTGS initiation threshold (Figure 4), indicating that almost every locus produce abRNAs, at least at some levels, and confirming that local and/or transient passing of the PTGS initiation threshold is sufficient to initiate S-PTGS, which can become systemic providing that the level of mRNA target is above the systemic S-PTGS threshold.

Whereas transcription strictly conditions S-PTGS amplification but only partly conditions S-PTGS initiation, another regulatory layer only conditions S-PTGS amplification. This layer relies on the capacity of DCL2-dependent 22-nt sRNAs to trigger the production of secondary siRNA by addressing targeted RNAs to RDR6 (Chen et al. 2010; Cuperus et al. 2010) whereas PTGS execution (*i.e.* RNA cleavage) mostly relies on DCL4-dependent 21-nt sRNAs. The reduced level of systemic S-PTGS observed in *dcl2* mutants (Parent et al. 2015; Taochy et al. 2017) implies that the production of 22-nt siRNAs is limiting. *DCL2* and *DCL4* transcripts levels appear similar at the organ level in vegetative tissues, which in part excludes the hypothesis that the limited production of 22-nt siRNAs is due to a disequilibrate ratio of DCL4/DCL2 (Liu et al. 2009). Instead, it is assumed that DCL2 and DCL4 do not process dsRNAs equally, with DCL4 being much more active than DCL2 (Parent et al. 2015; Taochy et al. 2017). Supporting this hypothesis, increased S-PTGS was observed in *dcl4* mutants (Parent et al. 2015; Taochy et al. 2017). By limiting S-PTGS, DCL2’s obscuration by DCL4 also contributes to protecting endogenous genes against the risk of undergoing systemic S-PTGS in cases of accidental RQC failure occurring in certain cells of the plants. Indeed, previous work showed that when essential RQC components are constitutively dysfunctional, endogenous genes produce siRNAs, leading to the plant’s death, and that plant growth can be rescued by mutating S-PTGS components such as *DCL2, RDR6* or *SGS3* (Martínez de Alba et al. 2015; Zhang et al. 2015; Scheer et al. 2021; Krzyszton and Kufel 2022; Feng et al. 2024). Moreover, associating *dcl4* with mutants of non-crucial RQC actors such as *xrn4* and *ski2* leads to a synthetic lethality phenotype largely attributed to PTGS of *NIAs* genes promoted by 22-nt siRNAs (Wu et al. 2020). These findings support the idea that DCL4 prevents highly expressed endogenous genes to amplify S-PTGS in conditions propitious to S-PTGS initiation, i.e. when RQC is impaired. However, it has never been shown that endogenous genes could undergo systemic S-PTGS in situations where RQC is functional and where the level of target mRNA is not artificially increased using a *Pro35S*-driven transgene. Nevertheless, we previously reported that a non-transgenic tobacco *nia* mutant in which metabolic derepression allows increasing the level of the endogenous *NIA* mRNAs to the level observed in *Pro35S:NIA2* transgenic plants become capable of undergoing systemic PTGS upon grafting (Palauqui and Vaucheret 1995). This prompted us to assess whether endogenous *NIA* genes expressed at physiological level could undergo systemic S-PTGS in scions grafted onto a rootstock producing *NIA* siRNAs and if DCL4 was playing a role in this process. Wildtype Arabidopsis plants grafted onto *Pro35S:NIA2* rootstocks did not undergo S-PTGS whereas *dcl4* mutants showed systemic S-PTGS upon grafting (Figure 5), indicating that the amount of target *NIA* mRNAs in wildtype plants is insufficient to sustain systemic S-PTGS as long as DCL4 is present to limit the production of 22-nt siRNAs by DCL2.

In conclusion, we propose a model (Figure 6 and Supplemental Figure 6) wherein two distinct thresholds condition S-PTGS initiation and amplification, the former being related to abRNAs targeted by RQC and the later to target mRNAs. The latter is directly conditioned by the transcription rate, whereas the former may depend also on other factors, *e.g.* the genomic environment and/or the arrangement of the DNA. According to this model, when the quantity of abRNAs produced by a locus saturates the RQC machinery, S-PTGS is initiated, sometimes in a very limited number of cells. The siRNAs produced in these cells target homologous mRNAs. If the amount of target transcripts is low, every mRNA is degraded and no secondary siRNAs are made because DCL4 obscures DCL2. Despite the possible movement of DCL4-derived 21-nt siRNAs to adjacent cells, S-PTGS does not become systemic because of the consumption of these siRNAs (Devers et al. 2020). In contrast, if the amount of target transcripts is high, part of them can be used to produce secondary siRNAs both in the initiating cells and subsequently in recipient cells in which primary siRNAs move, leading to systemic S-PTGS. Therefore, systemic S-PTGS can only occur if: i) RQC is locally and/or transiently saturated by abRNAs, and ii) if a sufficient amount of target mRNAs is present in every cell. This model applies to transgenes and endogenous genes. However, the evolvement of an efficient RQC machinery and of DCL4 obscuring DCL2 explains why S-PTGS is naturally never observed on endogenous genes, except in the cases of non-essential genes that underwent duplication (Coen and Carpenter 1988; Clough et al. 2004; Tuteja et al. 2004, 2009; Della Vedova et al. 2005). It also explains why natural accessions lacking DCL4 activity have never been reported. Indeed, only DCL2’s obscuration by DCL4 could allow RQC to dysfunction locally without translating into the drama of systemic S-PTGS. In contrast, transgenes expressed under strong promoters of viral origins have a high propensity to trigger systemic S-PTGS, and this likely relates to S-PTGS evolving first as a defence against viruses producing extremely high amounts of both conventional and non-conventional RNAs that can saturate RQC.

**Figure 6:**
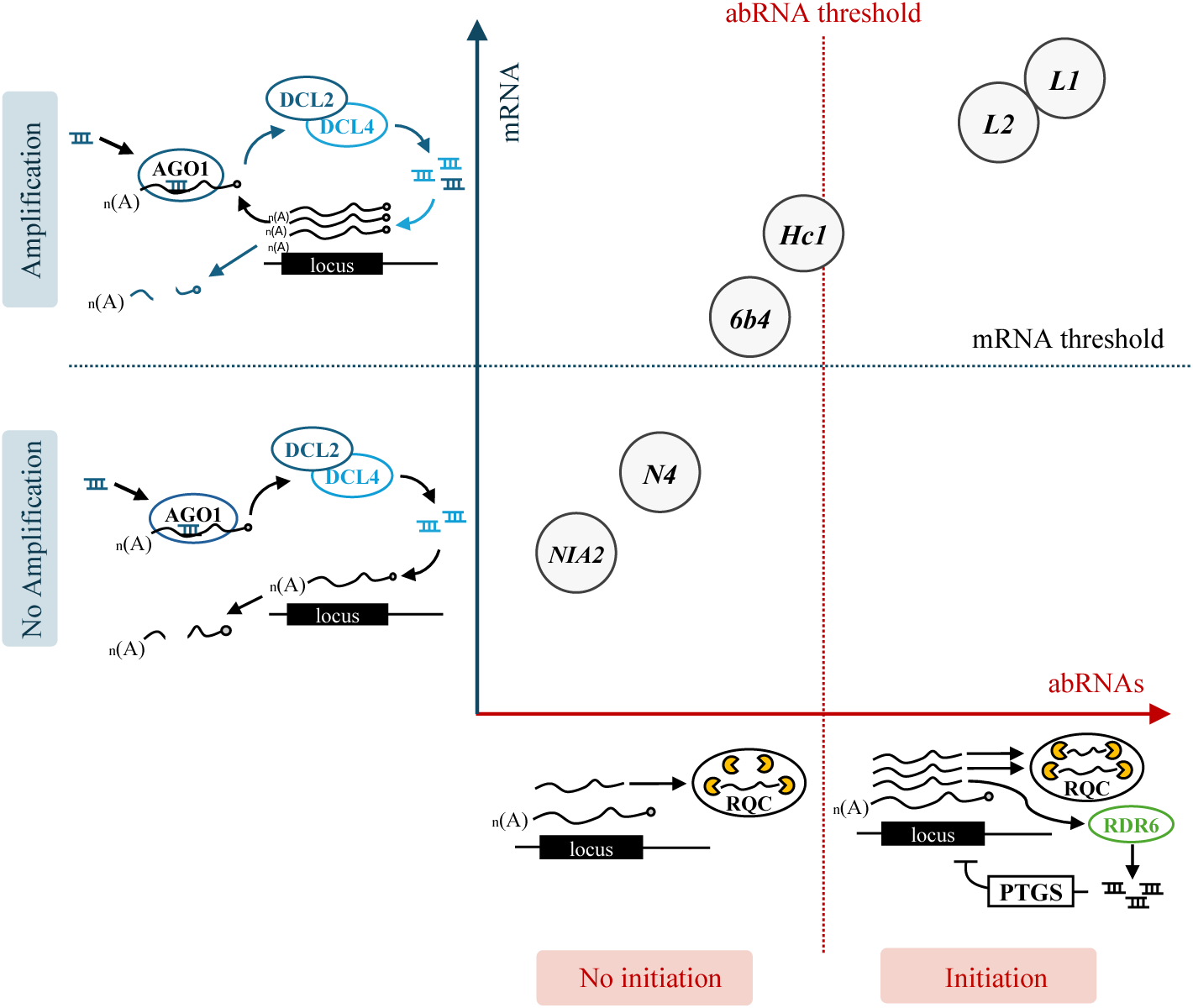
Model for two distinct thresholds conditioning S-PTGS initiation and amplification. S-PTGS is initiated when the quantity of abRNAs produced by a locus passes the initiation threshold and saturates the RQC machinery. It is conditioned by the transcription rate, but likely depends also on other factors, *e.g.* the genomic environment and/or the arrangement of the DNA. This initiation step likely happens in a very limited number of cells. The siRNAs produced in these cells target homologous mRNAs. If the amount of target transcripts is low, every mRNA is degraded and no secondary siRNAs are made because DCL4 obscures DCL2. Despite the possible movement of DCL4-derived 21-nt siRNAs to adjacent cells, S-PTGS does not become systemic because of the consumption of these siRNAs. In contrast, if the amount of target transcripts is high, i.e. above the amplification threshold, part of them can be used to produce secondary siRNAs both in the initiating cells and subsequently in recipient cells in which primary siRNAs move, leading to systemic S-PTGS. This model applies to transgenes and endogenous genes, which can be classified in three categories: i) those that are below the initiation and amplification thresholds and never trigger S-PTGS spontaneously but only when impairing both RQC and DCL4. A representative of this category is the endogenous *NIA2* gene, ii) those that are below the initiation threshold but above the amplification threshold. They never trigger S-PTGS spontaneously but only when grafted onto a silenced plants or when brought into an RQC-deficient mutant background. A representative of this category is the transgene locus *6b4*, iii) those that are above the initiation and amplification thresholds and trigger systemic S-PTGS. A representative of this category is the transgene locus *L1*.

## Methods

### Plant material and transformation

All Arabidopsis plants used in this study are of the Columbia (Col-0) accession. The transgenic lines *Pro35S:GUS* (*L1*, *L2*, *Hc1*, and *6b4*) (Elmayan et al. 1998; Béclin et al. 2002) and *ProSUC2::hpPDS* (*JAP3*) (Smith et al. 2007) as well as the *ProSUC2:GUS* line (Brioudes et al. 2021) have been described previously. The mutant lines *ago4-3* (Havecker et al. 2010)*, ago4-*5 (Greenberg et al. 2011)*, cmt3-*7 (Lindroth et al. 2001)*, dcl3-3* (Smith et al. 2007), *dcl4-2^gabi^*(Xie et al. 2005), *dcl4-2^E583K^* (Yoshikawa et al. 2005), *dcl4-5* (Deleris et al. 2006), *drm2-3* (Naumann et al. 2011)*, jmj14-4* (Le Masson et al. 2012), *mom1-2* (Moissiard et al. 2014), and *sgs2-1* (*rdr6*) (Mourrain et al. 2000) and *sgs3-1* (Mourrain et al. 2000) were also previously characterized.

The different *Pro35S:GUS* lines were generated by transforming Col-0 plants with the *Pro35S:GUS-tRbcS/tNOS-NPTII:ProNOS* construct via *Agrobacterium tumefaciens*-mediated transformation (Clough and Bent, 1998). Transformants were selected on kanamycin, and the number of T-DNA loci was subsequently determined by analyzing segregation ratios of their progeny on kanamycin-containing medium

The *iGUS* line is a monolocus transgenic line obtained by *Agrobacterium tumefaciens*-mediated transformation of the *Pro35S:GUS* line *6b4* with the construct *iGUS* (see the cloning and vector part). The initial T1 plants have been selected based on seeds red-fluorescence under a fluorescent microscope (Nikon® SMZ1500 Stereomicroscope).

### Growth conditions

For photography, seeds were directly sown in soil and grown in a greenhouse under long-day conditions. For all other experiments, seeds were surface-sterilized and sown on plates containing a nutrient medium (1.3% S-medium, Duchefa; 1% Phytoblend agar). Plants were vernalized at 4°C for two days and then transferred to a growth chamber set at 23°C with 70% humidity and a light intensity of 120 µE·m⁻²·s⁻¹. Plants were grown under either long-day (16 h light/8 h dark) or short-day (8 h light/16 h dark) conditions. After two weeks, seedlings were either harvested or transferred to soil and continued to grow under the same day-length conditions.

### EMS mutagenesis

For mutagenesis, approximately 1300 *L1 JAP3 jmj14-4* seeds were incubated in 5 mL of a 0.3% ethyl methanesulfonate (EMS) solution for 16 hours. To neutralize the EMS, 5 mL of a 1 M sodium thiosulfate (Na_2_S_2_O_3_) solution was added to the tube for 5 minutes. The seeds were then washed three times with water for 20 minutes each, suspended in 0.1% agarose, and stratified at 4°C for two days before being sown directly in soil. Seeds harvested from a bulk of three to five plants were screened for leaves with white nerves.

### Grafting techniques

The grafting protocol has been described previously (Turnbull et al. 2002). 6 days after germination seedlings were cut transversely across the hypocotyl with a razor blade. Scions and rootstocks were placed on a nitrocellulose filter (Hybond) and were introduced into a silicon microtube (2 mm long) to connect them to each other. Then, grafted plants were grown under short-day conditions for 7 to 14 days. Seedlings that did not show adventitious roots were transferred to soil and grown under a short-days photoperiod.

### RNA extraction, RT-qPCR and northern blot analysis

High and low molecular weight RNAs were extracted from 14 days after germination seedlings as previously described (Mallory et al., 2009). Northern blots for mRNAs and siRNAs were performed using 10 µg of RNA as previously described (Gy et al., 2007). *GUS* m-/si-RNAs, *PDS* siRNA, *U6* snRNA and *25S* rRNA probes have been previously described (Butel et al., 2017; Le Masson et al., 2012; Smith et al., 2007). The probe used for *NIA* siRNAs revelation corresponds to the 3’ region of *NIA2* (see Supplemental Table 1). Hybridization signals were revealed with a Typhoon FLA 9500 imager. The *25S* rRNA and *U6* snRNA probes were used to normalize the RNA loading for mRNAs and siRNAs, respectively.

### Methylation sensitive qPCR

Shoots from 14 days after germination seedlings were harvested and ground using a mortar and pestle. DNA was extracted from 100mg of the resulting powder using the NucleoSpin Plant II kit (Macherey-Nagel: 740770.50). Approximately 100 ng of DNA was digested with one of the following methylation-sensitive restriction enzymes: MspI (ThermoFisher: ER0541), HpaII (ThermoFisher: ER0511), or HaeIII (NEB: R0108S). 100 ng of undigested DNA was used as a control. qPCR was performed using the SsoAdvanced Universal SYBR Green Supermix kit (Bio-Rad: 1725274). The 35S-F2 (TGAGACTTTTCAACAAAGGG) and 35S rev primers (AAGGATAGTGGGATTGTGCG) were used for the amplification (Le Masson et al. 2012; Butel et al. 2021). The percentage of methylation was quantified by comparing the digested samples to the non-digested control.

### GUS activity measurement by fluorometric assay

Total proteins were extracted by grounding individual seedling or leaves pieces in a phosphate buffer (50 mM Na_2_HPO_4_, 50 mM NaH_2_PO_4_, pH7, 10 mM EDTA) with beads. Protein concentration in plant crude extract was quantified using the Bradford assay (Biorad) (Bradford 1976). GUS activity was measured by adding in excess GUS substrate (2 mM 4-methylumbelliferyl-b-D-glucuronide, Duchefa) to approximately 0.5µg of total protein. Accumulation of the fluorescent GUS reaction product (4-methylumbelliferon) was assessed by measuring fluorescence (excitation 365nm, emission 455 nm) with a fluorometer (Thermo Scientific Fluoroskan Ascent®) during 30 minutes with a period of 20 seconds. Maximal GUS activity in each sample (expressed as arbitrary unit of fluorescence by minute, U Fluo.min^−1^) was normalized by the protein concentration in the corresponding sample (U Fluo.min^−1^.µg of protein).

### Cloning and vector constructs

The construct *iGUS* (β-estradiol inducible *GUS* construct) and the *Pro35S:GUS* used to produce the new population of transformants was obtained using the GoldenBraid (GB) cloning system (Sarrion-Perdigones et al., 2011). All assembly reactions were made into the binary plant expression vector *pDGB3*. The different parts, their characteristics, origin and sequences are shown in Supplemental Table2. Reactions were performed as recommended by the GoldenBraid website (https://goldenbraidpro.com/https://goldenbraidpro.com/search/features/) with the T4 DNA ligase (ThermoFisher: EL0011) and the restriction enzymes Esp3I (BsmBI, ThermoFisher: ER0451) and BsaI (NEB: R3733S). All parts GB parts used in this study are described in the Supplemental Table2.

For the *iGUS* construct, we adapted a pre-existing β-estradiol inducible system to the GoldenBraid cloning system. The different parts of the system were domesticated from the *pMDC7* vector and cloned in individual *pUPD2* vectors (*ProG1090*, *XVE*, *ProLexA*). The CDS of the chimeric transcription factors *XVE* was cloned in a *pDGB3*-alpha1 vector downstream of the plant constitutive *G1090* promoter (*ProG1090*) (Ishige et al., 1999) and upstream of the *HSP18.2* terminator (*tHSP18*.2) (Nagaya et al., 2010) domesticated from *A. thaliana* genome. The *GUS* CDS was assembled with the inducible *LexA* promoter (*ProLexA*) and the *tRbcSE9* in a *pDGB3*-alpha 1 vector, and a *DsRed* transcriptional unit under the control of the *ProCsVMV* and the *tNOS* was assembled into a *pDGB3-*alpha2 vector (plant selection marker). While the alpha1 *ProG1090:XVE-tHSP18.2* unit was switched to a *pDGB3-*omega1 vector with the help of an alpha2 twister plasmid containing a stuffer fragment (*SF*), the alpha1 *ProLexA:GUS-tRbcSE9* and alpha2 *ProCsVMV:DsRed-tNOS* units were assembled in a *pDGB3-*omega2 destination plasmid. Finally, the omega1 *ProG1090:XVE-tHSP18.2/SF* vector was combined with the omega2 *ProLexA:GUS-tRbcSE9/ProCsVMV:DsRed-tNOS* into a *pDGB3*-alpha2 destination vector to obtain the *iGUS* vector (*ProG1090:XVE-tHSP18.2/SF/ProLexA:GUS-tRbcSE9/ProCsVMV:DsRed-tNOS*).

For the *Pro35S:GUS* construct, the GUS transcriptional unit was assembled in a *pDGB3-alpha1* vector (*Pro35S:GUS-tRbcSE09*) and the resistance cassette into a pDGB3-alpha2 (*ProNOS:NPTII-tNOS*). Then, both transcriptional units were assembled into a *pDGB3-omega2* to produce the final *Pro35S:GUS* vector (*Pro35S:GUS-tRbcSE09*/ *ProNOS:NPTII-tNOS*).

The resulting *iGUS* and *Pro35S:GUS* vectors were transferred into *A. tumefaciens* (strain *C58pmp90*) and used to transform *A. thaliana* plants by floral dip (Clough and Bent 1998).

## Acknowledgments

We thank Taline Elmayan and IJPB members for fruitfull discussions and the IJPB greenhouse keepers for taking care of the plants. We also thank Elodie Akary, Lionel Gissot and Martine Pastuglia who provided some of the GoldenBraid modules used in this study.

## Author contributions

M.L, N.B., I.L.M. and H.V conceived and designed the study. All authors performed the experiments. M.L, N.B., A.B., I.L.M. and H.V analyzed the data. H.V wrote the manuscript with contributions from M.L., N.B. and A.B.

---

**Supplemental Figure 1:**
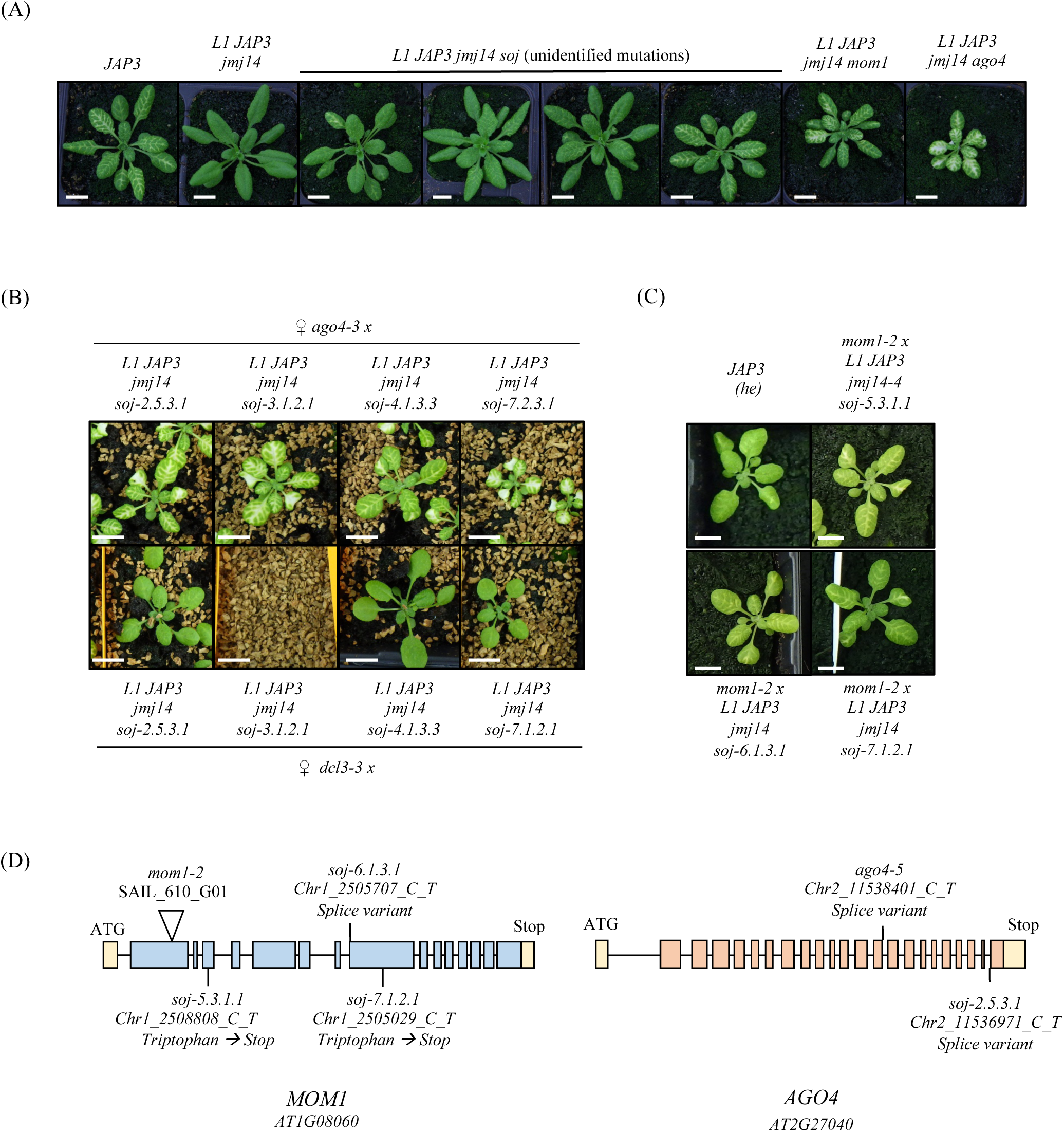
Mutations in *AGO4* and *MOM1* suppress the effect of *jmj14* on *JAP3* silencing. (A) Range of phenotypes of suppressor mutants isolated from the mutagenesis of the *L1 JAP3 jmj14* line. *L1 JAP3 jmj14 ago4* and *L1 JAP3 jmj14 mom1* exhibit the higher levels of photobleaching. Scale bars, 1cm. (B, C) Complementation assay of the different *L1 JAP3 jmj14 ago4* (B) and *L1 JAP3 jmj14 mom1* (C) mutants. Control crosses with *dcl3* mutants (B) or heterozygous *JAP3* lines (C) shows the silenced *JAP3* phenotype. Scale bars, 1cm. (D) Schematic representation of *AGO4* and *MOM1* including the localization of the different *soj* mutations identified in this study. The filled squares correspond to exons.

**Supplemental Figure 2:**
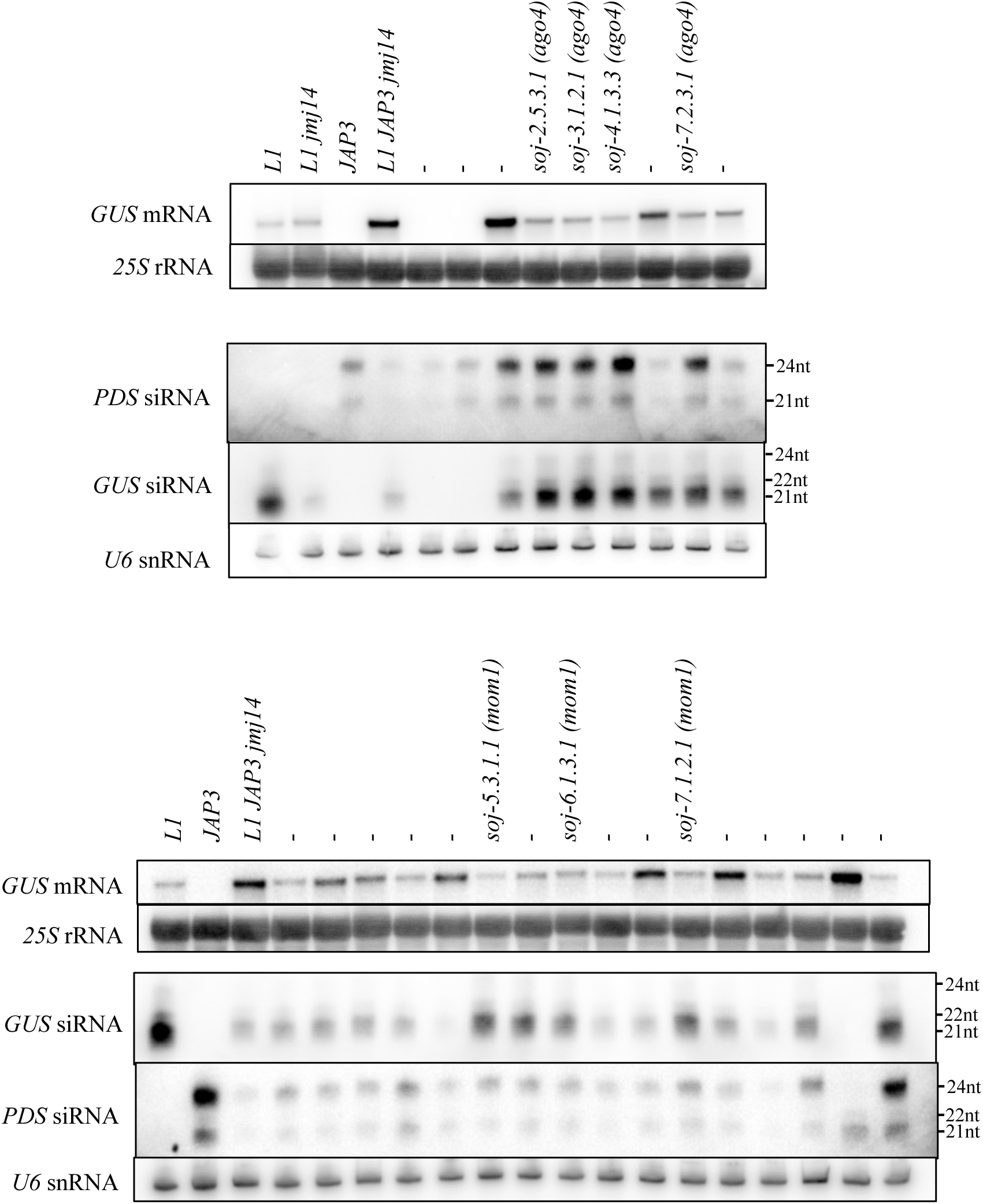
Original blots of the Figure 2.

**Supplemental Figure 3 :**
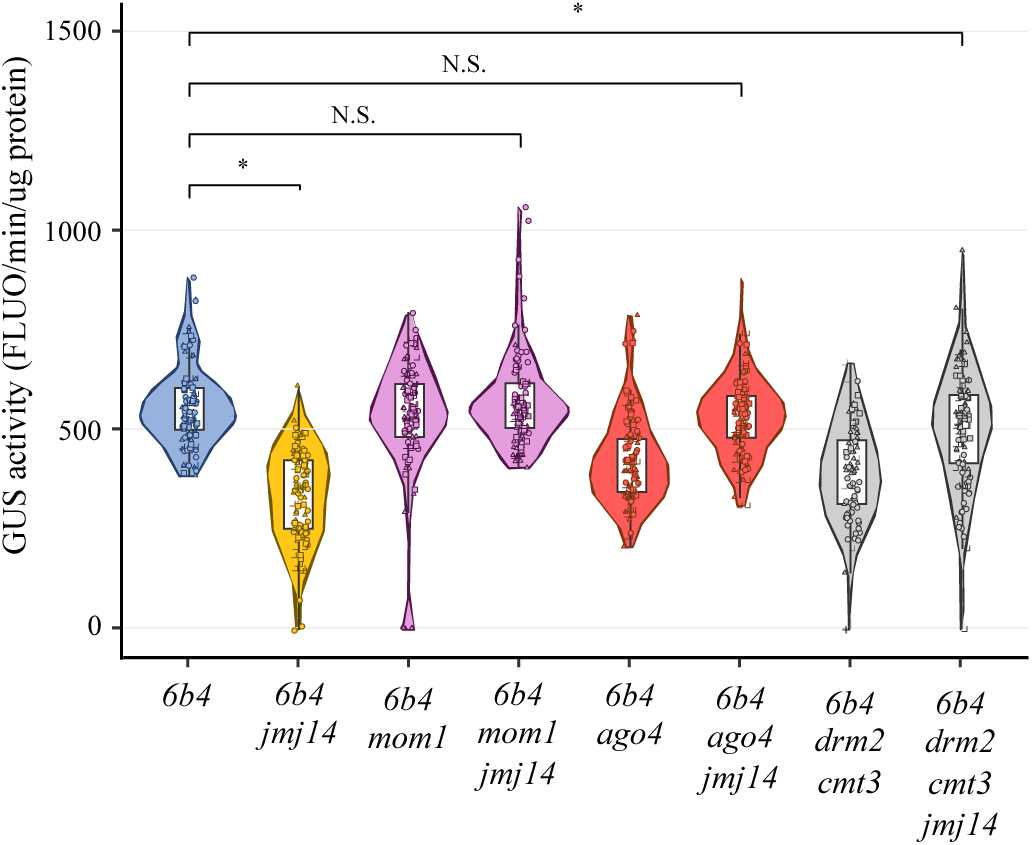
Mutations in *AGO4, MOM1* and *DRM2/CMT3* restore *6b4* GUS activity at wildtype levels in a *JMJ14*-deficient background. GUS activity in aerial parts of 14 days after germination measured in *6b4* and *6b4 jmj14* plants carrying second site mutations (*ago4*, *mom1* and *drm2 cmt3*) suppressing the effect of *jmj14*. Each dot represents an individual plant, with more than 80 plants tested per genotype. P-values were calculated using non parametric Wilcoxon tests adjusted with Holm–Bonferroni correction (* means P<0.05 and N.S., not significant).

**Supplemental Figure 4 :**
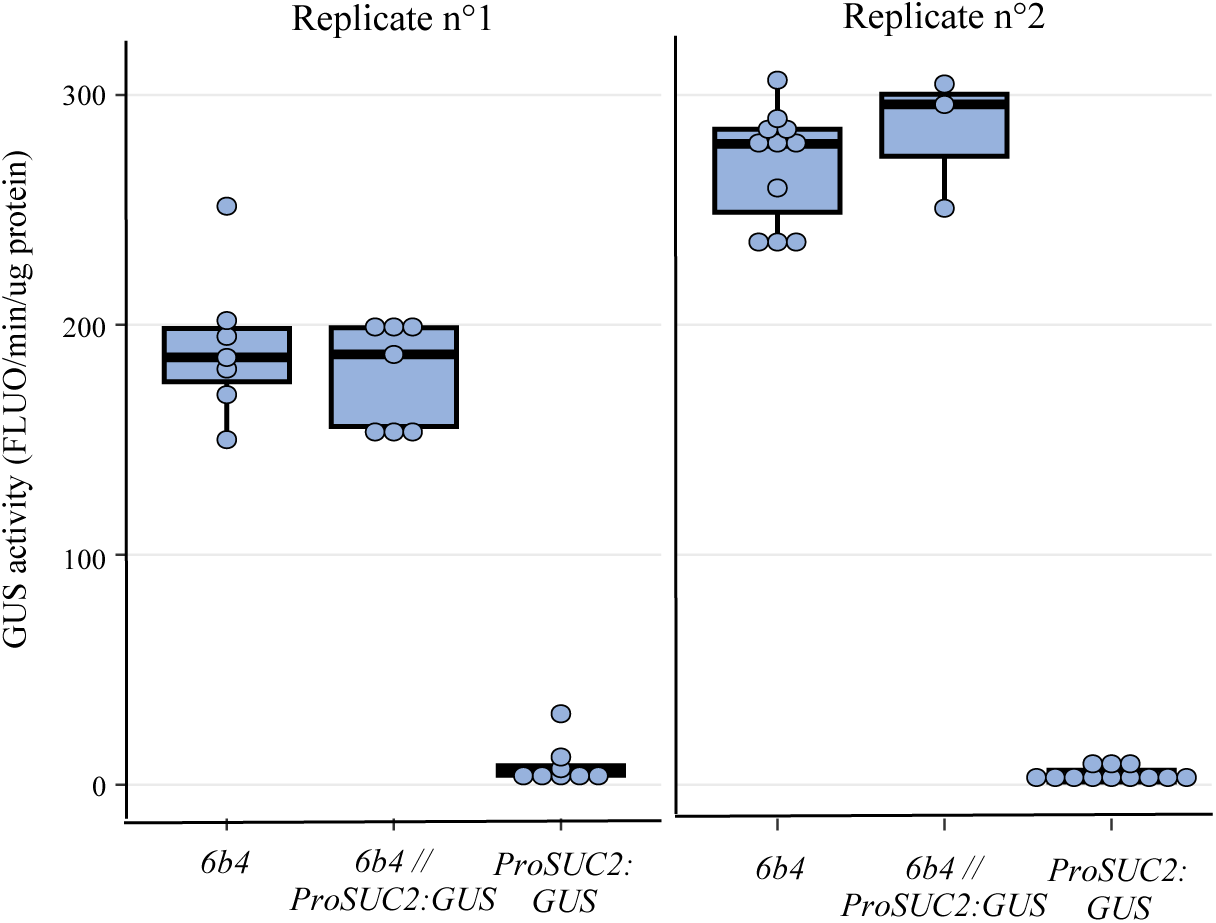
The *ProSUC2:GUS* line does not produce a systemic silencing signal. GUS activity was measured in leaves of *6b4*, *ProSUC2:GUS* and *6b4* plants grafted onto the *ProSUC2:GUS* line (*6b4 // ProSUC2:GUS*). Two independent grafting experiments are shown as separated panels.

**Supplemental Figure 5 :**
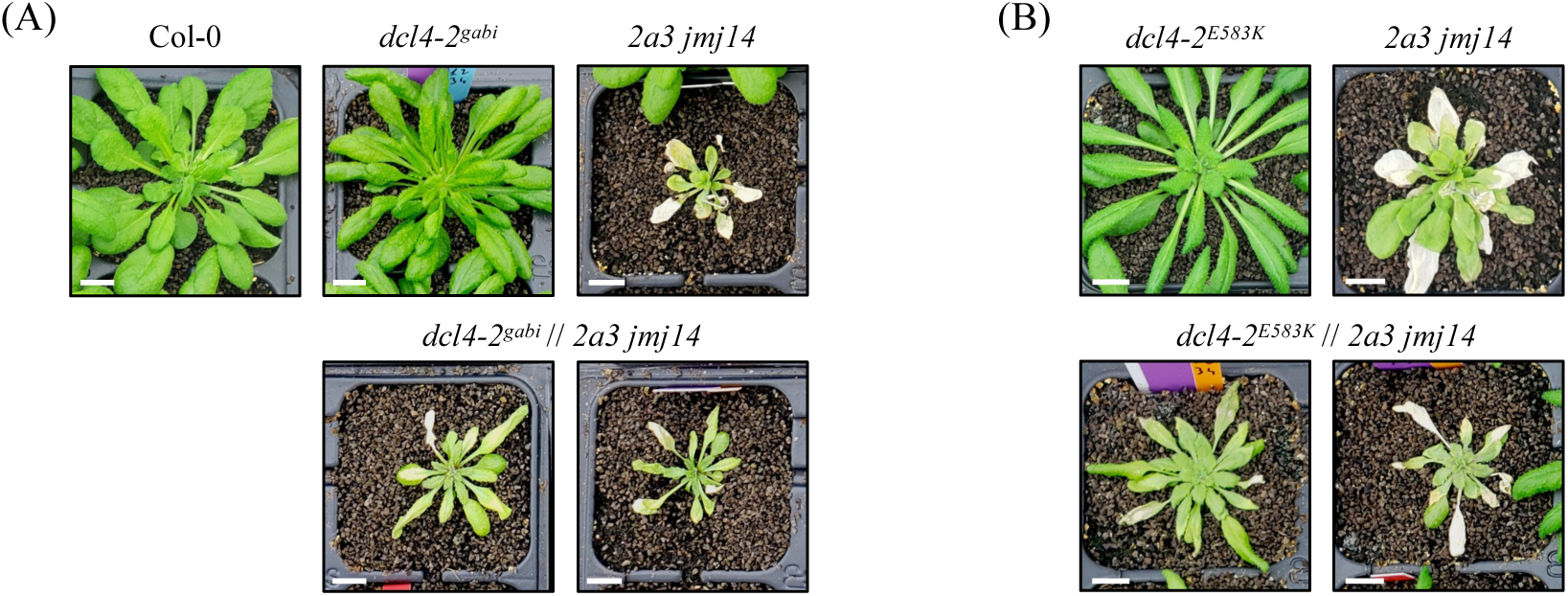
All *dcl4* alleles tested allow endogenous *NIAs* genes to trigger systemic PTGS upon grafting onto the *NIA* siRNA producing rootstock *2a3 jmj14*. Images of aerial parts of distinct *dcl4* alleles grown on their own roots or grafted onto a silenced *Pro35S:NIA2* transgenic line (*2a3*) in a *jmj14* mutant background. (A) *dcl4-2^gabi^*. (B) *dcl4-2^E583K^*. *dcl4-5* is shown on Figure 5. Scale bars, 1cm.

**Supplemental Figure 6 :**
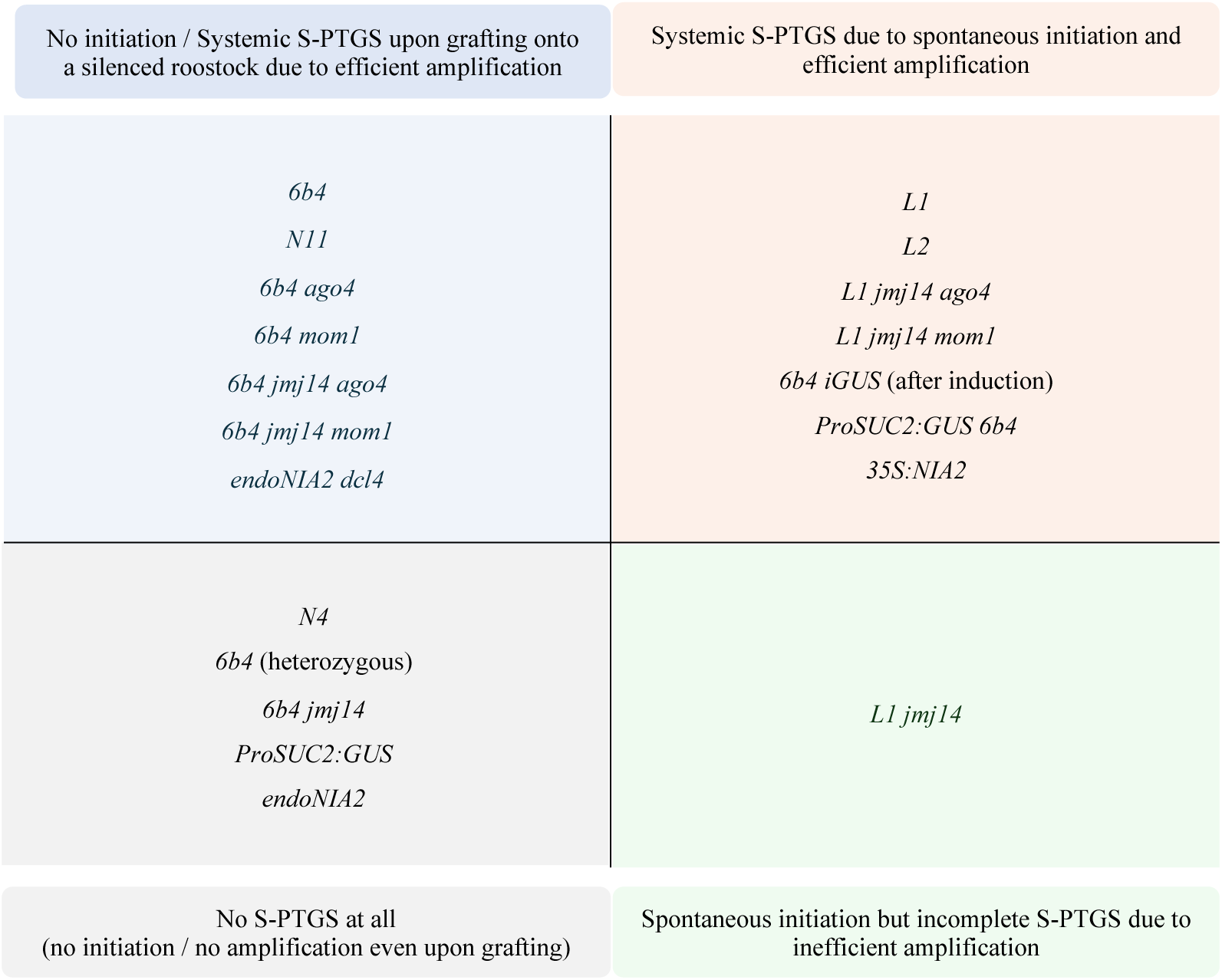
Initiation and amplification status of the different lines used in this study.

